# Feasibility of high-resolution perfusion imaging using Arterial Spin Labelling MRI at 3 Tesla

**DOI:** 10.1101/2023.08.02.551576

**Authors:** Sriranga Kashyap, ĺcaro Agenor Ferreira Oliveira, Kâmil Uludağ

## Abstract

Cerebral blood flow (CBF) is a critical physiological parameter of brain health and it can be non-invasively measured with arterial spin labelling (ASL) MRI. In this study, we evaluated and optimised whole-brain, high-resolution ASL as an alternative to the low-resolution ASL employed in the routine assessment of CBF in both healthy participants and patients. Two high-resolution protocols (i.e., pCASL and FAIR-Q2TIPS (PASL) with 2 mm isotropic voxels) were compared to a default clinical pCASL protocol (3.4 × 3.4 × 4 *mm*^3^), all of whom had an acquistion time of ≈ 5 min. We assessed the impact of high-resolution acquisition on reducing partial voluming and improving sensitivity to the perfusion signal, and evaluated the effectiveness of z-deblurring on the ASL data. We compared the quality of whole-brain ASL acquired using three available head coils with differing number of receive channels (i.e., 20, 32 and 64 ch). We found using higher coil counts (32 and 64 ch coils as compared to 20 ch) offer improved SNR and improved acceleration capabilities that are beneficial for ASL imaging at 3T. The inherent reduction in partial voluming effects with higher resolution acquisitions improves the resolving power of perfusion without impacting sensitivity. In conclusion, our results suggest that high-resolution ASL (2 - 2.5 mm isotropic) has the potential to become a new standard for perfusion imaging at 3 T and increase its adoption into clinical research and cognitive neuroscience applications.

## Introduction

Arterial spin labelling (ASL) is a non-invasive neuroimaging technique that uses magnetically labeled arterial blood water as an endogenous tracer to measure Cerebral Blood Flow (CBF)^1, 2^. ASL provides a safe and repeatable method for assessing brain state and function without any risk of toxicity or allergic reactions from exogenous contrast agents. ASL can also be utilised to assess quantitative CBF in units of mL/100g/min at an individual voxel level^1, 3^.

In recent years, technological advances in MRI scanner hardware and software and new cutting-edge analysis methods have positively impacted the range of ASL applications and resulted in notable growth in the number of publications^4–6^. Another factor for its increasing popularity in clinical research is the community effort to standardise acquisition methods, data structures, and analyses^7–9^. However, widely adopted standards (e.g., described in the ASL ‘white paper’^7^) prescribe spatial resolutions of 3-4 mm in-plane and 4-8 mm slice thickness for ASL scans that are maximally 5-6 minutes long (typical length of clinical research / standard-of-care MRI protocols) but may not be optimal anymore with current hardware and MRI sequences. While these protocols may suffice for macroscopic effects (such as pattern of large regions of hypo-perfusion), they are insufficient to detect subtle abnormalities that may represent early-stage of neurological diseases or small lesions^10^. Therefore, ASL at higher spatial resolution (*<* 3 mm nominal, isotropic) is highly desirable.

Another reason for going to high spatial resolutions is to reduce partial volume (PV) effects, which occur when the voxel signal contains fractional contributions from more than one tissue type, e.g., grey matter (GM), white matter (WM) and cerebrospinal fluid (CSF). This can introduce inaccuracies in perfusion quantification of the tissue of interest resulting in either or both under-estimation and over-estimation, depending on the PV fractions in the voxels. For instance, Asllani et al.^11^ show that a voxel mixture of 80:20% grey:white matter (this ratio would be inclusive after threshold, in most cases), would result in a 24% perfusion under-estimation. Another example is the study^12^, investigating the impact of a higher-resolution ASL protocol compared to low-resolution Positron Emission Tomography (PET) scans, and they demonstrate that uncorrected CBF PET images might underestimate the grey matter (GM) CBF by 20%. In fact, in 2006, Donahue et al.^12^ actually envision the future of ASL imaging at 3 T to be higher resolutions of 2.5mm in-plane or higher. 17 years since, 3 T ASL imaging is still routinely carried out with voxel sizes *>* 3 mm, and the voxels are almost never isotropic, which can lead to underestimation of lesions and even misdiagnosis in the direction of the lowest spatial resolution. While there haven been methods and algorithms developed that can provide a means to post-hoc correct for PV effects^13–15^, they can usually not recreate lost information and, therefore, the most straight-forward and preferred approach is to just acquire the data at higher spatial resolutions.

This is notwithstanding high-resolution ASL studies carried out at field strengths higher (4.7 T, 7 T) than that typically used in the clinic (1.5 T, 3 T). For example, Mora Álvarez et al.^10^ demonstrated the feasibility of a high-resolution Continuous ASL (CASL) at 4.7 T within a clinical time frame of 6 minutes.

The study also observed reduced PV averaging at 1.5 × 1.5 × 3*mm*^3^ resolution. Another interesting example is the study published by Zuo et al.^16^ where they employed Turbo-FLASH (Fast Low Angle SHot) ASL, both pseudo-continuous ASL (pCASL) and pulsed ASL (PASL) at 7 T showing the feasibility of achieving an in-plane resolution of 0.85 × 1.7*mm*^2^. At 7 T, recent functional MRI (fMRI) studies also showed the feasibility of using perfusion-weighted contrast with ASL at sub-millimetre spatial resolutions of 0.9 mm isotropic^17^ and 0.7 mm isotropic^18, 19^ using a 3D-EPI^20^ readout with a FAIR^21^ QUIPSS II^22, 23^ labelling scheme.

While there is evidence of the transformative potential that ultra-high field scanners can have for clinical research and cognitive neuroscience applications, they are limited in availability compared to the ubiquity of 3 T scanners. Therefore, a translation of high-resolution ASL to widely available 3 T clinical platforms is urgently needed to catalyse clinical research as well as further advance the standards-of-care. This requires systematic optimisation attuned to easily accessible workflows, currently not explored in existing ASL literature.

The current study addresses these aforementioned challenges and gaps in literature by first, developing, testing, and evaluating high-resolution ASL protocols at 3 T in clinically feasible times and compare them to a vendor default protocol that are typically used in routine clinical scanning. To this end, we developed optimised 2 mm isotropic pCASL and PASL protocols that balances the trade-off between signal-to-noise (SNR) and acquisition time (TA) to be feasible for clinical application (TA ≈ 5 minutes). Furthermore, we also evaluated the impact of the the choice of standard head coils on 3 T perfusion imaging. Therefore, we systematically evaluated our protocols and the clinical default protocol with all three commercially available head coils (20 channel head & neck, 32 channel head only, 64 channel head & neck) to ascertain the optimal hardware for high-resolution acquisitions. Additionally, we quantify and demonstrate the reduction in partial voluming enabled by the high-resolution acquisitions. Finally, the lengthening of the readout with 3D-GRASE is recognised to result in through-plane (z-axis) blurring resulting in loss of spatial resolution^24, 25^. We also assess the impact of advanced post-processing methods such as z-deblurring to improve spatial fidelity of the acquired data.

## Methods

### Participants

Eight healthy volunteers (4 females and 4 males, mean age = 29 ± 4 years) participated in the study and provided written informed consent prior to scanning. All participants were screened healthy individuals, non-smokers, not taking any medications, and with no history of neurological or neurovascular conditions. All procedures in this study conformed to the standards set by the Declaration of Helsinki and was approved by the Research Ethics Board of University Health Network according to the guidelines of Health Canada.

### Data acquisition

Data were acquired on a Siemens Magnetom Prisma 3 T MRI scanner (Siemens Healthineers, Erlangen, Germany) at the Slaight Family Centre for Advanced MRI (Toronto Western Hospital, Toronto ON, Canada) having a maximum gradient strength of 80 mT/m, slew rate of 200 T/m/s and running on the XA30A IDEA software platform. We used three commercial MRI coils, namely, a 20ch Head and Neck, a 32ch Head only, and a 64ch Head and Neck for receive, and the transmit was carried out by the body coil. Participants were positioned by taking the eye centres as reference for the magnet isocentering to minimise B_0_ offsets for the labelling in the neck. All data in participant were acquired in the same scan session. The participants were brought out of the scanner, coils were exchanged, and the participants were repositioned to the magnet’s isocentre. The sequential order of coils was pseudo-randomised between participants to avoid any systematic biases.

#### Anatomical imaging

Structural scans were acquired with the 32ch Head coil. Whole-brain anatomical data were acquired using a 3D Multi-Echo Magnetisation-Prepared Rapid Gradient Echo (3D-MEMPRAGE) sequence^26^ that uses volumetric EPI navigators combined with selective data reacquisition^27^ to produce (prospectively) motion-corrected T_1_w images^28^ that were used in the study. 3D-MEMPRAGE data were acquired at 0.8 mm isotropic resolution, TI = 1000 ms, TEs_1 − 4_ = 1.81, 3.6, 5.39, 7.18 ms, TR = 2500 ms, *α* = 8^*º*^, 208 sagittal slices, matrix = 320 × 320, GRAPPA = 2, Ref. lines = 32, partial-Fourier_*slice*_ = 6/8, echo spacing = 11.2 ms, bandwidth = 740 Hz/pixel, turbo factor = 168, total acquisition time ≈ 8 min). The four echoes were combined (using root mean squares, RMS) into a high fidelity T_1_-weighted image following the scanner’s on-line reconstruction. Quantitative T_1_ mapping was carried out using a 3D Magnetisation-Prepared 2 Rapid Gradient Echoes (3D-MP2RAGE) sequence^29^. The MP2RAGE T_1_ maps were only used to facilitate perfusion quantification and thus were acquired at a 1.2 mm isotropic resolution, TIs_1 − 2_ = 700, 2500 ms, *α*_1 − 2_ = 4^*º*^, 5^*º*^, TE = 4.04 ms, TR = 3200 ms, 144 axial slices, matrix = 192 × 192, GRAPPA = 2, Ref. lines = 32, partial-Fourier_*phase*_ = 6/8, echo spacing = 9.08 ms, bandwidth = 150 Hz/px, turbo factor = 144, total acquisition time ≈ 4 min). T_1_ maps were calculated in-line using the Siemens MapIt package (Siemens Healthineers, Erlangen, Germany).

#### Perfusion imaging

All ASL protocols were developed using the Siemens Advanced 3D-ASL work-in-progress (WIP) sequence (courtesy of Siemens Healthineers, Erlangen, Germany) available for the XA30A baseline platform. The ASL data were acquired with a segmented 3D-GRASE readout for improved SNR^30–32^. Three ASL protocols were acquired per coil in each participant: 1) the clinical default protocol (3.4 × 3.4 × 4 mm, ‘Clinical’ in Table 1); 2) a high-resolution (or hires) pCASL protocol (2 mm isotropic, ‘Hires’ in Table 1); and 3) a hires PASL protocol employing a FAIR Q2TIPS^33^ labelling scheme (2 mm isotropic, ‘PASL Hires’ in Table 1). For clinical and hires ASL variants, two steady-state magnetisation (M_0_) calibration images were acquired without any labelling, but with matched readout and TR increased to 20 s, one of the M_0_ had the opposite phase-encoding for distortion-correction. The new hires protocols developed in this study were acquired in approximately the same total time as the spatially anisotropic clinical ASL scan (≈ 5*min*).

**Table 1.**
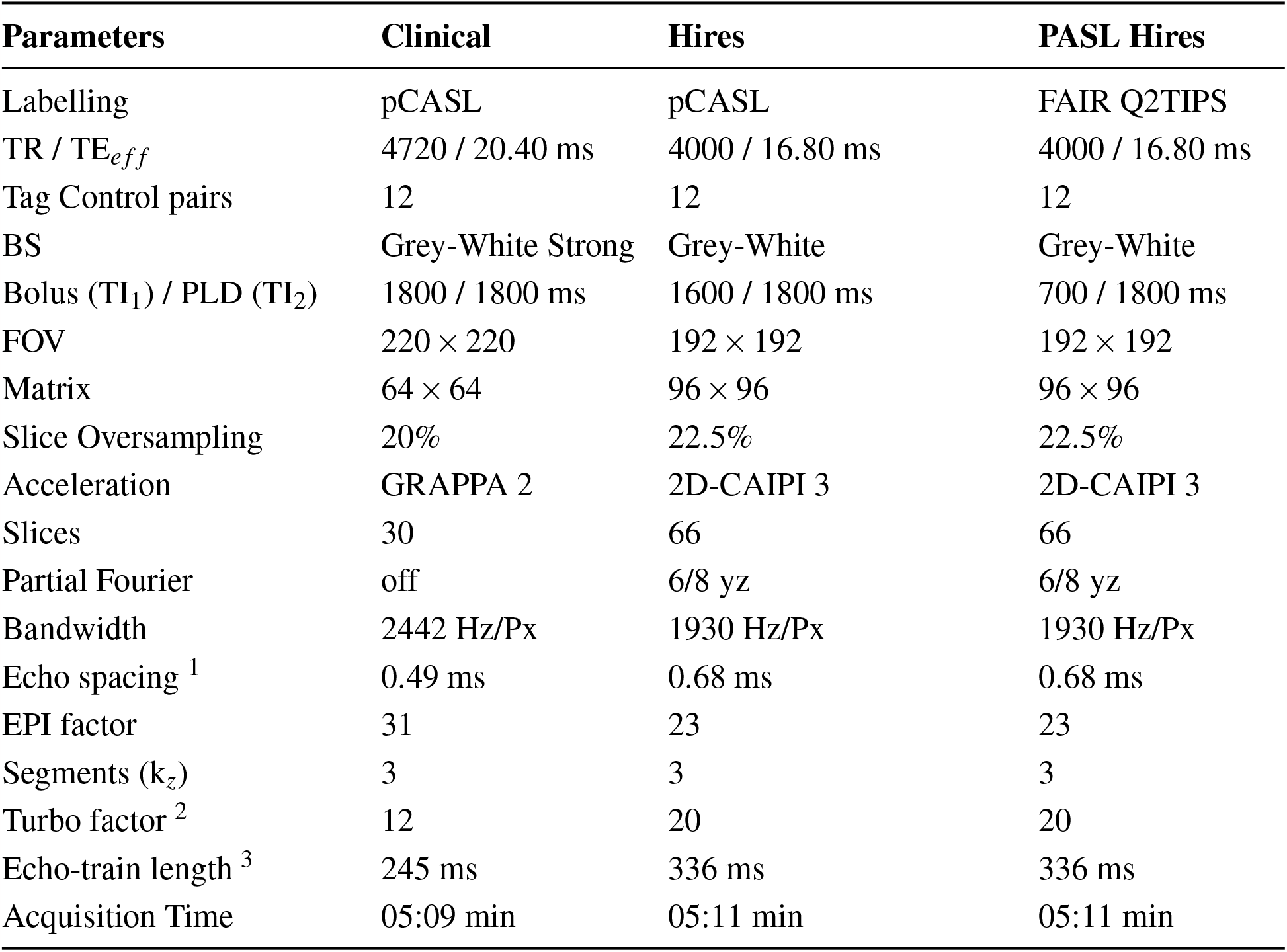
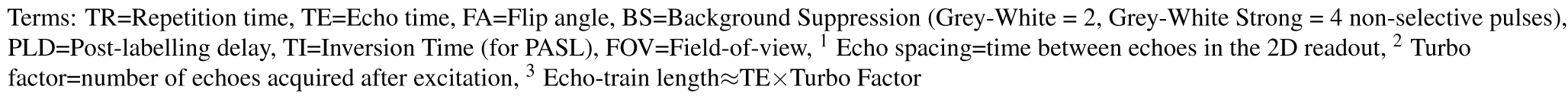
Table of sequence parameters for the three ASL protocols in the present study. Although the study focuses on the two pCASL protocols, the PASL protocol is included here for completeness.

**Table 2.** Table of FWHM (in mm) estimated using AFNI’s *3dFWHMx* for the clinical and hires 3D-GRASE datasets deblurred using the FFT method and direct kernel estimation as implemented in *oxasl*_*deblur*. Comparison of FWHM for different deblurring methods can be found in Table S2 (Supplementary Material). Numerical values presented are mean ± std. dev across participants.

| FWHM | Clinical | Clinical Deblurred | Hires | Hires Deblurred |
| --- | --- | --- | --- | --- |
| x | $5.51 \pm 0.27$ | $5.49 \pm 0.76$ | $3.08 \pm 0.18$ | $2.65 \pm 0.21$ |
| y | $6.24 \pm 0.20$ | $5.91 \pm 0.55$ | $2.85 \pm 0.17$ | $2.42 \pm 0.22$ |
| z | $8.73 \pm 0.65$ | $5.22 \pm 0.49$ | $4.01 \pm 0.31$ | $1.41 \pm 0.12$ |
| ACF | $10.08 \pm 0.67$ | $7.24 \pm 0.58$ | $4.83 \pm 0.36$ | $2.69 \pm 0.12$ |

### Data Processing

#### Anatomical imaging

The RMS-combined, motion-corrected T_1_-weighted 3D-MEMPRAGE was processed using Freesurfer v 7.3.2^34–36^(https://surfer.nmr.mgh.harvard.edu/) using a brain mask that was generated using *mri_synthstrip*^37^ and was provided as an additional input to the *recon-all* pipeline.

#### Perfusion imaging

The first volume of the ASL timeseries was discarded as separate M_0_ scans had been acquired for quantification. The pre-processing steps were carried out using FSL^38^ included motion and distortion-correction, where all control and label volumes were independently realigned to the first volume of the ASL scan. The separately acquired M_0_ scans were rigidly registered to the first volume of the ASL scan and then distortion correction was done using FSL’s *topup*^39^ with the two M_0_ images. The perfusion timeseries were calculated using sinc-subtraction as implemented in FSL’s *perfusion_subtract*. The M_0_ images, perfusion-weighted data, and the MP2RAGE T_1_ maps (coregistered to M_0_) were used as input to *oxasl*^40^ (https://github.com/physimals/oxasl) for voxelwise perfusion quantification. The M_0_s were co-registered to the anatomical image using Freesurfer’s *bbregister*^41^ to obtain CBF maps in both native and structural space. No adaptive spatial smoothing^42^ or partial-volume correction^43^ were applied. Next, all anatomical scans were carefully registered to the 1 mm isotropic MNI Non-linear 2009c asymmetric template space^44, 45^ using ANTs SyN algorithm^46, 47^ (https://github.com/ANTsX/ANTs). Native space maps from *oxasl* were resampled in a single step to the MNI space using *antsApplyTransforms*. The stability of the perfusion signal over time (temporal SNR, tSNR) was calculated dividing the temporal mean by the temporal standard deviation of the perfusion-weighted data (also referred to as, perfusion tSNR). The SNR (consequently, tSNR) of a voxel is expected to scale proportionally with its volume and thus, makes it challenging to compare datasets of highly different spatial resolutions. Therefore, to better appreciate the tSNR relative to a dataset’s spatial resolution, the perfusion tSNR map from the hires scan was scaled by the ratio of the voxel volumes of clinical to hires datasets (46.24 mm^3^/ 8 mm^3^ = 5.78).

#### Partial volume analysis

In order to visualise the impact of the higher spatial resolution acquisition, participant-wise T_1_-weighted images were resampled to the nominal spatial resolution of the clinical protocol (3.4 × 3.4 × 4.0 mm^3^) or the hires protocol (2.0 mm isotropic). The resampled T_1_-weighted images were segmented using FSL’s *fsl_anat*^48^ to obtain PV estimates. The cortical grey matter segmentation from Freesurfer was morphologically dilated by 1 voxel and resampled to the two resolutions and this resampled, dilated cortical mask was used as the ROI for the PV analyses. To this end, we used a histogram-based analysis to first sort the voxels into different PV fraction bins. Then, to compare the two different acquisition resolutions, the number of voxels in each histogram bin was scaled by their voxel volumes of 46.24 mm^3^ and 8 mm^3^, respectively, for the clinical and hires protocols giving us the volume of PV voxels in each bin. This normalisation enabled direct comparison of the PV. A difference between the hires and clinical histograms (after rescaling) were computed for all values above a PV fraction threshold of 0.5 for each of the three tissue classes, namely, GM, WM and CSF.

#### Deblurring analysis

In an additional analysis, ASL data acquired from the 32ch coil were pre-processed using the *oxasl*_*deblur* (https://github.com/physimals/oxasl_deblur). We evaluated two different methods for deblurring the data, namely, fast Fourier Tranform division (FFT) and Lucy-Richardson deconvolution (Lucy) as implemented in *oxasl*_*deblur* with three different kernel options (direct estimation, lorentzian, lorentzian with a weiner filter). Smoothness of the deblurred data was estimated using AFNI’s^49, 50^ *3dFWHMx*^51^ function.

## Results

### Comparison of clinical and hires ASL data

Figure 1 shows the group average absolute CBF (in units of mL / 100g / min) maps from clinical and hires pCASL protocols presented in three orthogonal views (middle panel) for the three head-coils used to acquire the data (drawing in left panel). The panel on the right shows the distribution of the CBF values in GM across the participants’ data as a violin plot with the CBF values represented on the y-axis for the two protocols. The figure annotation represents the mean ± standard deviation of the distribution. A comparison of the group average perfusion-weighting and relative CBF (rCBF, in arbitrary units) for the two protocols and three head-coils is shown in Supplementary Figure S1. It is important to note that the rCBF calculated using *oxasl* is the perfusion-weighted image (PWI) following kinetic model inversion (i.e., one step before M_0_ calibration to physiological units of mL / 100g /min) and is not relative to the whole brain mean or normal white matter. For the clinical and hires protocols, we observe that the mean CBF values are very similar for all three coils. The CBF values obtained from the clinical protocols are ≈ 17% greater than those obtained from the hires data and both measures are in the acceptable range for healthy volunteers^7^. Summary statistics for all the different perfusion metrics calculated from the data are tabulated in Supplementary Tables S2-S6.

**Figure 1.**
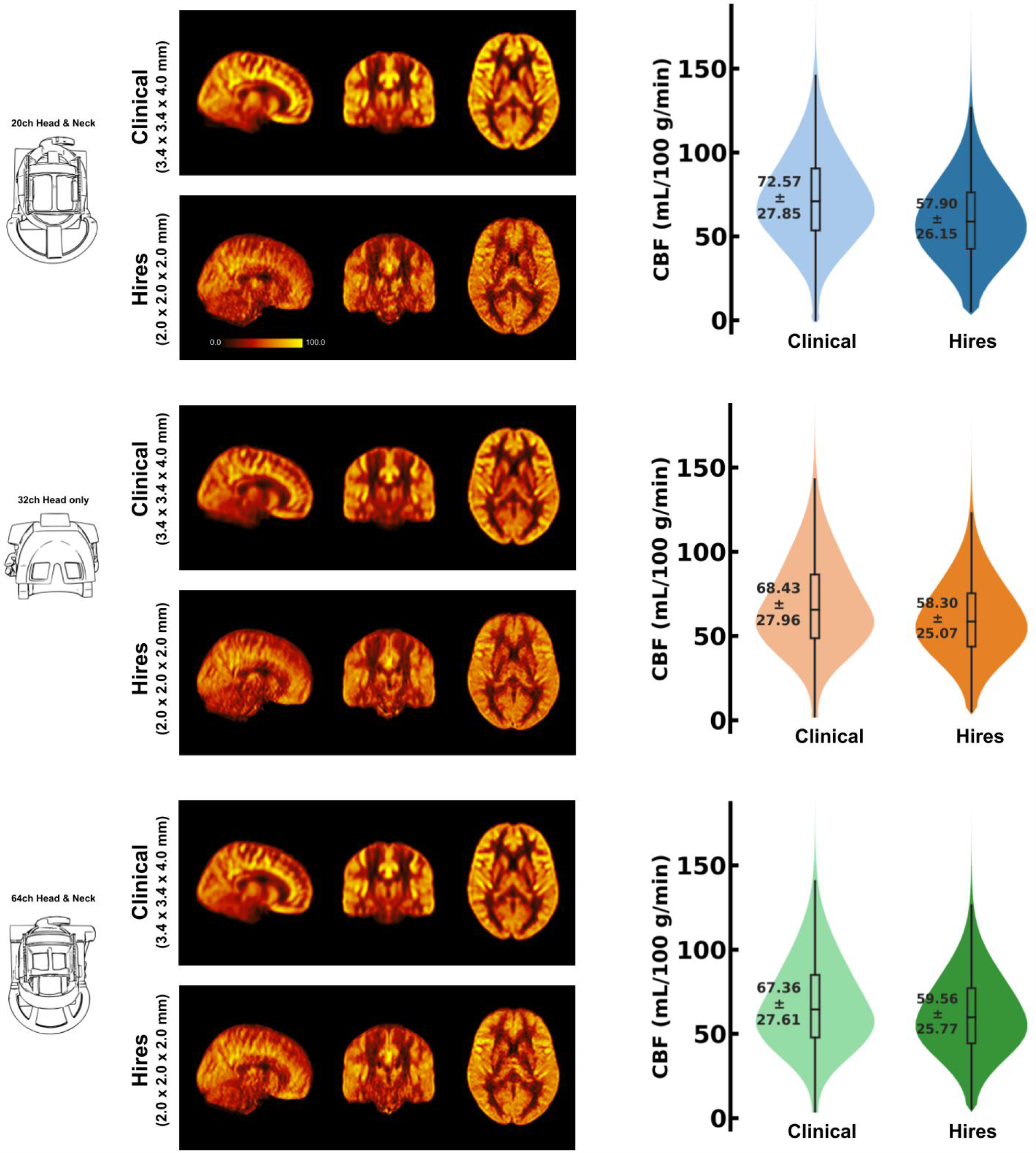
Mosaic of orthogonal views of the group-average (n=8) CBF (in mL / 100g / min) for data acquired using the three respective head coils (drawing on the left). In the middle column, the maps obtained from the clinical and hires pCASL acquisitions are displayed in the top and bottom rows, respectively. (Right) Violin plots of the CBF distribution across all participants’ data (n=8) for the two acquisitions. The annotation represents the mean ± standard deviation of the distribution.

### Analysis of the partial voluming

One of the primary advantages of acquiring higher spatial resolution data is to reduce partial voluming of the signal of interest. In Figure 2, the difference in voxel volumes (hires−clinical) are plotted at each partial volume fraction bin ranging from 0.5-1.0 (50% to ‘pure’ single tissue composition) for three tissue classes, that is, GM (a), WM (b) and CSF (c) using a dilated GM ROI. Data from each participant is shown as a coloured dot, with the mean across participants plotted as a black dashed line. In Figure 2a, we observe that on average, above a PV fraction of 0.6 (60% GM), there is a net positive change in the volume of GM and remains positive for all higher PV fractions. In other words, even within the dilated GM ROI, there is a larger volume (total ≈ 6713 mm^3^) of ‘pure’ GM in the hires compared to the clinical data, therefore, partial voluming is reduced. This finding is corroborated by the spatial maps of PV as illustrated in the right panels with the PV map of hires and clinical shown on top and bottom rows, respectively. A similar pattern is observed for Figure 2b and 2c that quantifies the PV in WM and CSF respectively. That is, within the dilated GM ROI used to extract these results, there is a significantly larger volume of ‘pure’ WM (total ≈ 16938 mm^3^) and ‘pure’ CSF (total ≈ 15422 mm^3^). As GM is bound on either side with WM and CSF, we can infer that greater the number of ‘pure’ non-GM voxels, lower is the amount of voxels which are PV with GM, and this finding is corroborated by the spatial maps of PV.

**Figure 2.**
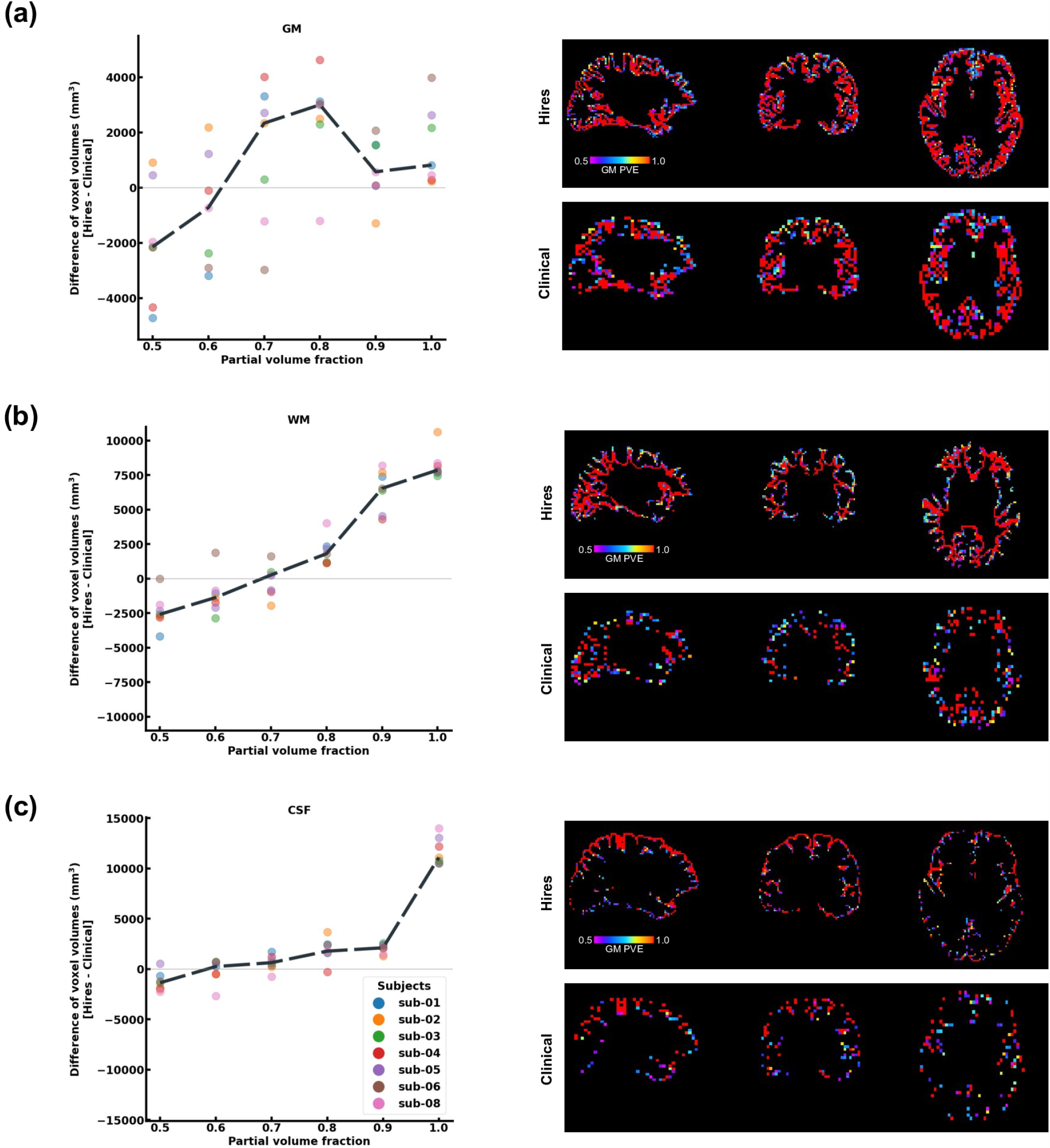
Histogram difference plot for all bins ≥ 0.5 threshold of PV fraction for (a) GM, (b) WM, (c) CSF tissue classes of the hires and clinical acquisitions. Value from each participant is represented as a colour-coded circle and the group average is plotted as a black dotted line. A single-participant PV estimate map is shown in the right panel for the clinical and hires spatial scales, spatially illustrating the findings of the histogram-analysis. Please note, sub-07 is excluded from this analysis as *f sl*_*anat* could not be completed.

### Deblurring analysis of 3D-GRASE ASL

Table 2 shows that the effective spatial resolution of both the clinical and hires datasets is different to what is indicated in the protocol, also referred to as the nominal spatial resolution (in this study, 3.4×3.4×4.0 mm^3^ and 2.0 mm isotropic respectively). Systematic evaluation of five parameter combinations in *oxasl*_*deblur* (Table S2) shows that all five combinations result in an improvement in a reduction in the Full Width at Half Maximum (FWHM). We found that using the FFT method with direct kernel estimation to yield the smallest effective FWHM (clinical: 6.38 ± 0.33 mm vs hires: 2.38 ± 0.13 mm, Table S2).

Table 2 shows the FWHM estimated from AFNI’s 3*dFWHMx* for x-,y- and z-axes as well as the effective FWHM (ACF). We observe that irrespective of the acquisition resolution, the smoothness is maximal along the z-axis (Clinical: 8.73 ± 0.65 mm, Hires: 4.01 ± 0.31 mm) and this is the axis along which *oxasl*_*deblur* is most effective, reducing the smoothness estimate to 5.22 ± 0.49 mm and 1.41 ± 0.12 mm for Clinical and Hires data respectively. The change in the estimated FWHM along z after deblurring (ΔFWHM_*clinical*_/ΔFWHM_*hires*_) is 1.35 times larger for the hires dataset than the clinical data. Figure 3 shows the group-average CBF maps for the clinical and hires datasets before (‘orig’) and after (‘deblurred’) deblurring and the distribution of CBF values across all participants’ data is also shown as a violin plot.

**Figure 3.**
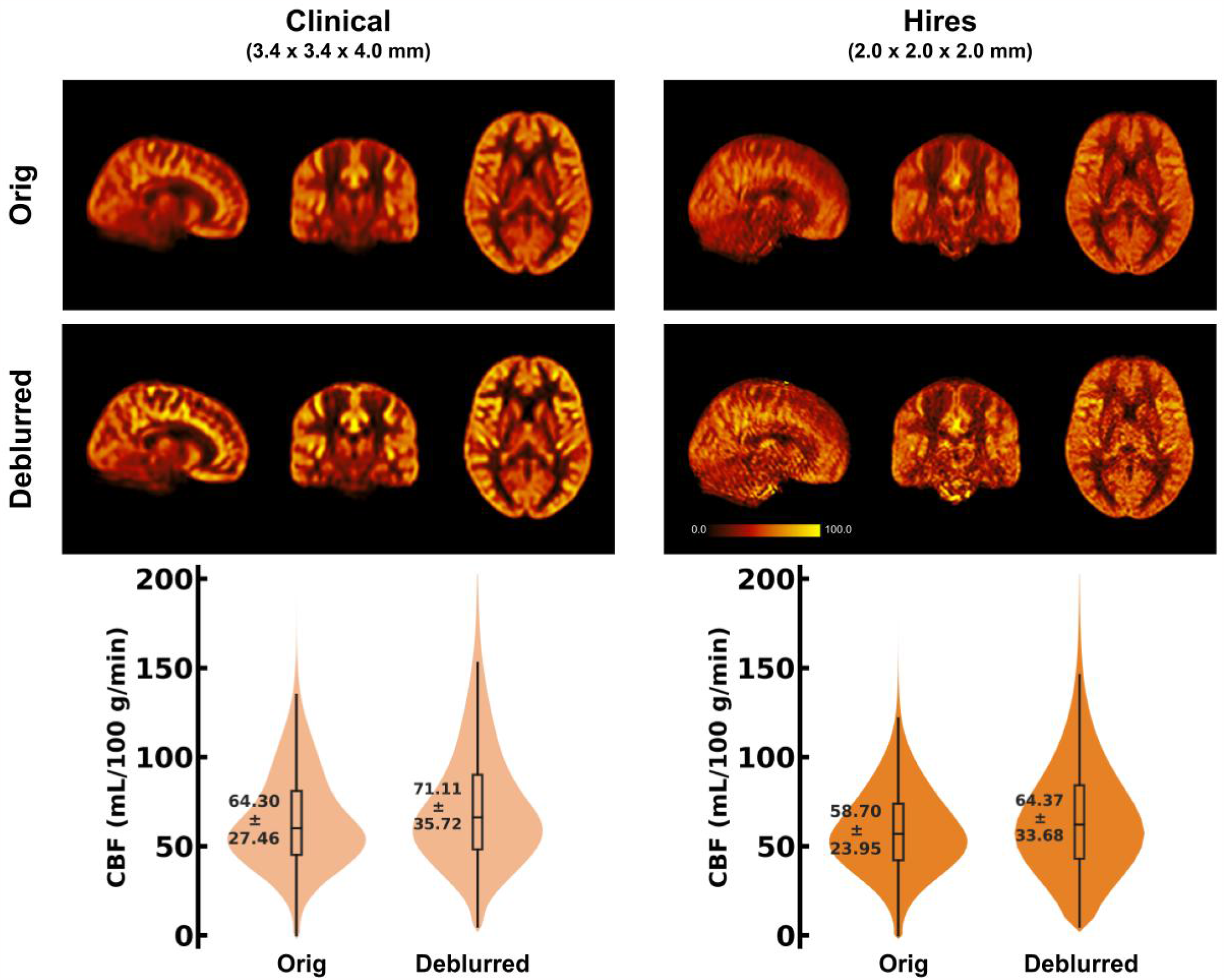
(Top) Orthogonal views of the group-average CBF maps (in mL / 100 g / min) (n=8) obtained using the 32 channel head coil, before and after deblurring using the FFT method and direct kernel estimation as implemented in *oxasl*_*deblur*. (Bottom) Violin plots of the CBF distribution across all participants’ data (n=8) for the two acquisitions before and after deblurring. The annotation represents the mean ± standard deviation of the distribution.

### Impact of head-coil choice for imaging perfusion

Figure 1 demonstrates that robust CBF maps can be acquired independently of the coil choice. However, the spatial distribution of the CBF maps from the hires protocol show a preference for the 32 and 64 ch. Figure 4 top and middle rows illustrate the impact of perfusion tSNR across the three coils. In the case of the clinical protocol, the increasing coil count has an ≈ 2-2.5% gain in perfusion tSNR, whereas, the hires protocol has an ≈ 34-42% gain in perfusion tSNR with increasing coil count (Table S5). The perfusion tSNR maps of the hires data, rescaled by the ratio of voxel volume (Figure 4, bottom row), illustrates the improvement of tSNR with 32 and 64 coils over the 20 ch. Additionally, Table S4 shows that the inter-quartile range (IQR) of the perfusion-weighting increases with increasing coil count (20/32/64 ch: for clinical, 312.50/334.30/338.79 a. u., and for hires 386.34/423.67/440.03 a. u.) for both protocols. The IQR of perfusion-weighting between the three coils behaves similarly with the hires PASL protocol (20/32/64 ch: 396.12/447.66/454.97 a. u., Table S9). Therefore, it is the SNR benefits afforded by higher coil count rather than quality or type of labelling used that is responsible of the improvement in the higher IQR of perfusion values.

**Figure 4.**
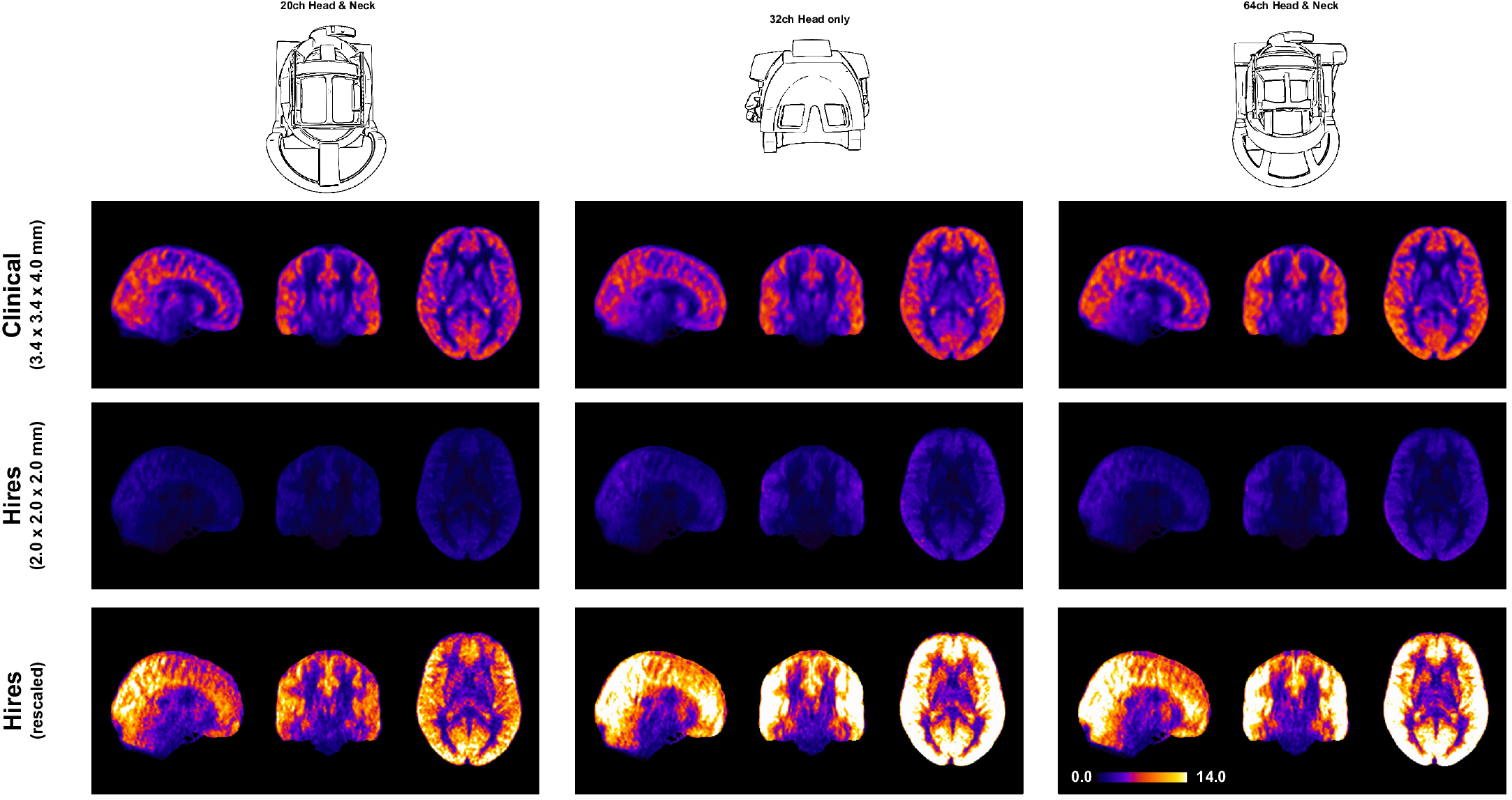
Orthogonal views of the group average perfusion tSNR maps (n=8) where the pCASL Clinical and Hires maps are presented in the top and middle rows respectively. The bottom row shows the hires tSNR data (middle row) but with rescaled values.

## Discussion

In this study, we demonstrate that it is feasible to measure perfusion robustly and repeatably using ASL at a high-spatial resolution of 2 mm isotropic within clinically-feasible times of ≈ 5 minutes. In this study, we used the updated version of the vendor provided 3D-GRASE ASL sequence (Siemens Advanced 3D-ASL WIP) and no custom sequence developments were carried out to enable wide-spread usage without the necessity to developing custom MR sequences or image reconstruction, despite the continuous progress being made on the development of ASL methods^8^). Therefore, we expect the sequence parameter choices made in this study can be selected in the 3D-GRASE ASL sequence available from the vendor on most modern scanners. We show that the increased spatial resolution does result in a reduction of partial voluming compared to the default clinical protocol. We show that through-plane blurring is a problem for 3D-GRASE ASL independent of the protocol being used. We find z-deblurring to be more effective on the hires than the clinical data. Finally, the choice of head coil for imaging perfusion with ASL at 3 T does play an important role with the 32 and 64 ch being particularly well-suited. Consistent with the results of deblurring, the hires datasets benefit most from perfusion tSNR improvements with higher coil counts.

### Impact of spatial resolution on ASL imaging

We show that increasing the spatial resolution of ASL 5.78× the clinical resolution does not have a detrimental effect on the measuring perfusion (Figure 1) and takes the same duration as a clinical scan (≈ 5 min). The mean perfusion-weighting values in the high-resolution data were found to be similar to the clinical data (e.g., 32-channel: 608.55 ± 256.24 vs. 605.91 ± 313.30 a. u.) (Table S4). Importantly, however, the hires perfusion-weighted images exhibited approximately 27% greater IQR (32-ch: clinical 334.3 vs. 423.67 a. u.) than the clinical data. Since the IQR is a measure of spread around the mean, this measure is indicative of the dynamic range of perfusion in the data. Being capable of resolving a wider range of perfusion values is critical to detect subtle abnormalities and early-detection of neurological diseases^52^, therefore, emphasising the importance of high-spatial resolution imaging^10^ in clinical research and cognitive neuroscience applications.

Supporting of the finding of an improvement in the dynamic range, acquiring data at a higher spatial resolution reduces PV effects. The cortical GM is bound on either side by WM and CSF and PV occurs when a GM voxel contains fractional distributions from these adjacent tissue classes that influence cortical perfusion measures. Figure 2 shows that hires ASL data consistently yields a greater volume of ‘pure’ tissue voxels than the clinical data (GM: ≈ 6713 mm^3^, WM: ≈ 16938 mm^3^, CSF: ≈ 15422 mm^3^) (Please note that these PV fractions were derived from a dilated, cortical GM ROI, that is, the ROI does not consist of the large ventricles or the majority of WM in the brain). The increased number of ‘pure’ WM and CSF voxels indicate that the hires data can enable a more effective separation of non-GM signal contributors to the perfusion signal of interest.

Partial volume correction was not performed at any stage of processing of the datasets^13, 43^. In the absence of PV correction of the lower resolution clinical protocol data, the lower CBF in WM partial voluming with GM would result in a reduction of the average CBF in GM. However, PV of GM with CSF (or rather vessels in CSF) can have the opposite effect resulting in higher than expected CBF values in GM, which is likely the case here. It is also important to note the default clinical protocol was not subject to any optimisation in the present work. While seemingly contrary to expectations, for parameter sets similar to the default clinical protocol, the CBF values in our data are consistent with studies that use a similar sequence^32^. Other reasons could be the fact that high-resolution acquisitions inherently reduce partial voluming effects and therefore, can be more sensitive to the CBF variability within GM. Maps including that of the perfusion-weighting and rCBF are shown in Figures S1 (clinical vs hires) and S2 (pCASL vs PASL). We found that the hires PASL results are in good agreement with the hires pCASL (Figure S2). Consistent with previous work^53, 54^, the pCASL labelling scheme exhibits about 22-26% higher perfusion tSNR than FAIR Q2TIPS for the hires acquisitions in our study.

### Impact of deblurring on 3D-GRASE ASL data

Due to high spatial resolution required, the total echo train length (TE * TF) can exceed 300 ms (*>> π×*T_2_^*∗*^ of tissue) resulting in increased blurring^55–57^, that occurs maximally in the slice direction (through-plane or z-axis). Thus, requiring post-processing correction or making compromises that would render whole-brain acquisitions infeasible. We find that application of z-deblurring has a demonstrable effect on the improvement of the spatial fidelity (or reducing the estimated FWHM) of the ASL data as shown in Table 2. Interesting to note that the FWHM along z for the deblurred clinical data (5.22 ± 0.49 mm) is still larger than the non-deblurred hires data (4.01 ± 0.31 mm). This has an important implication in clinical setting where advanced image post-processing is often unavailable. Importantly, the hires ASL protocols enable researchers and clinicians to resolve perfusion changes with a higher spatial fidelity (without requiring advanced image processing) than the post-hoc deblurred clinical datasets. Furthermore, post-processing deblurring methods have their limitations and they cannot synthesise resolution from information lost in acquisition. While lengthening the echo-train is an important concern, our findings (Table 2) indicate that deblurring methods are more effective for high-resolution ASL imaging.

### Impact of coil choice on ASL imaging

We demonstrate that robust relative CBF maps can be acquired independently of the coil choice (Figure 1), however, higher coil counts(32 and 64 ch), offer substantial gains in perfusion tSNR compared to the 20 ch coil (Figure 4). We find that increasing coil count has an ≈ 2-2.5% gain in perfusion tSNR for the clinical protocol compared to ≈ 34-42% gain for the hires protocol (Table S5). One reason for this difference could be that data acquired with clinical protocol in Figure 1 is relatively insensitive to the choice of coil due to its low spatial resolution (i.e., low thermal noise) and acceleration (i.e., no g-factor penalty) requirements. On the other hand, the hires protocols accelerate higher and have increased thermal noise relative the clinical protocol owing to the smaller voxel sizes, and therefore, benefits from the increased number of coils (Figure 4).

Interestingly, Figure 4 shows that the reducing the voxel size (i.e., higher spatial resolution) actually results in a gain in perfusion SNR (clinical vs hires (scaled)) which may seem counter-intuitive from the standpoint of conventional fMRI where the SNR of the BOLD signal decreases with increasing resolution. However, this is due to the different signal origins of the BOLD and perfusion contrasts. By reducing PV with veins and macrovasculature, we are reducing the signal contributors of the BOLD signal, whereas, these same signal components are sources of noise in perfusion imaging, as they have very low perfusion signals. In addition, reducing WM contribution of voxels dominated by GM improves the fidelity of GM perfusion values and reduced influence of physical noise stemming from WM. Therefore, reducing PV increases our sensitivity to the cortical microvasculature signal and recues noise and signal contribution from WM and CSF. In other words, higher spatial resolution decreases image SNR in both BOLD and perfusion methods due to reduction in the number of proton (i.e., voxel volume), but also reduces noise sources in perfusion imaging stemming from CSF, veins, and WM.

### Limitations

While we demonstrate clear benefits of high-resolution ASL imaging for clinical research and cognitive neuroscience applications (group studies), the present study is limited in its ability to comment on a potential impact in daily clinical practice (single subject, diagnostic). Nevertheless, we believe future studies investigating impact of ASL sequence parameters in routine clinical practice should employ a modestly higher isotropic resolution (e.g., 2.5 mm) to enable better visualisation of localised differences in perfusion. Here, we also opted for modest acceleration schemes (Table 1) as the protocols were to be compared on all three available head coils and the 20 ch coil would be the lowest common denominator. The availability of the 3D-GRASE readout with 2D-CAIPIRINHA undersampling enabled us to achieve higher isotropic spatial resolution for perfusion imaging^32, 58, 59^. For a future non-comparison type of study, this protocol optimisation can be pushed further, to take advantage of the higher coil count and achieve higher acceleration. A systematic exploration of different CAIPI acceleration schemes or trajectories, impact of reduced g-factor noise amplification on image quality is, unfortunately, beyond the scope of the present work.

### Concluding remarks

Taking together, this study demonstrates the feasibility and benefits of imaging perfusion using high-resolution isotropic ASL for clinical research and cognitive neuroscience applications at 3 T. We have shown that increasing the spatial resolution does not compromise accuracy, quality of the perfusion maps, and allows for a wider dynamic range of perfusion values. We show the high-resolution data can more effectively separate out the non-GM signal contributors (reduce PV effects) which improves sensitivity to cortical microvasculature and tissue in GM. Additionally, post-processing methods such as z-deblurring are important considerations for whole-brain perfusion imaging using 3D-GRASE ASL to improve the spatial fidelity of the data. High-resolution acquisitions take advantage of the higher coil counts and offer substantial gains in perfusion tSNR with the 32 and 64 ch coils. Echoing what Donahue and colleagues envisioned in 2006^12^, we strongly believe that high-resolution ASL (2 - 2.5 mm isotropic) can be a new standard for perfusion imaging using ASL at 3 T and be adopted into clinical and cognitive neuroscience research workflows.

## Data Availability Statement

The datasets will be made available via the Canadian Open Neuroscience Platform (https://portal.conp.ca/) [link upon publication].

## Author Contributions

SK: Conceputalisation, Methodology, Investigation, Data curation, Formal analysis, Visualization, Writing – original draft, Writing – review & editing; IAFO: Methodology, Investigation, Validation, Visualization, Writing – original draft, Writing – review & editing; KU: Conceputalisation, Methodology, Project administration, Funding acquisition, Resources, Supervision, Writing – original draft, Writing – review & editing.

## Acknowledgments

We thank Asma Naheed, MRT for the scanning support at the Slaight Family Centre for Advanced MRI, Toronto Western Hospital, University Health Network, Toronto, Canada. We like to thank Dr. Thomas F. Kirk for the invaluable discussions about the partial voluming problem in ASL MRI data. We like to thank Dr. Dimo Ivanov for his insights into ASL acquisition. We thank Dr. Gerald R. Moran and Dr. Josef Pfeuffer for their collaboration and providing the Advanced 3D ASL WIP package for the Siemens XA30 platform.

## Supplementary Material

### Impact of deblur options on the 3D-GRASE ASL data

**Table S1.** Table of FWHM (in mm) estimated using AFNI’s *3dFWHMx* for different deblurring methods available in *oxasl*_*deblur*. Numerical values presented are mean ± std. dev across participants. This evaluation was carried out on data from the 32 ch coil only.

|  | FFT |  |  | Lucy |  |
| --- | --- | --- | --- | --- | --- |
|  | Direct | Lorentz | Lorwien | Direct | Lorentz |
| Clinical | $6.38 \pm 0.33$ | $6.57 \pm 0.3$ | $6.65 \pm 0.33$ | $6.55 \pm 0.3$ | $6.55 \pm 0.35$ |
| Hires | $2.38 \pm 0.13$ | $2.53 \pm 0.12$ | $2.43 \pm 0.14$ | $2.54 \pm 0.09$ | $2.62 \pm 0.09$ |

### Comparison of clinical and hires pCASL protocols

**Figure S1.**
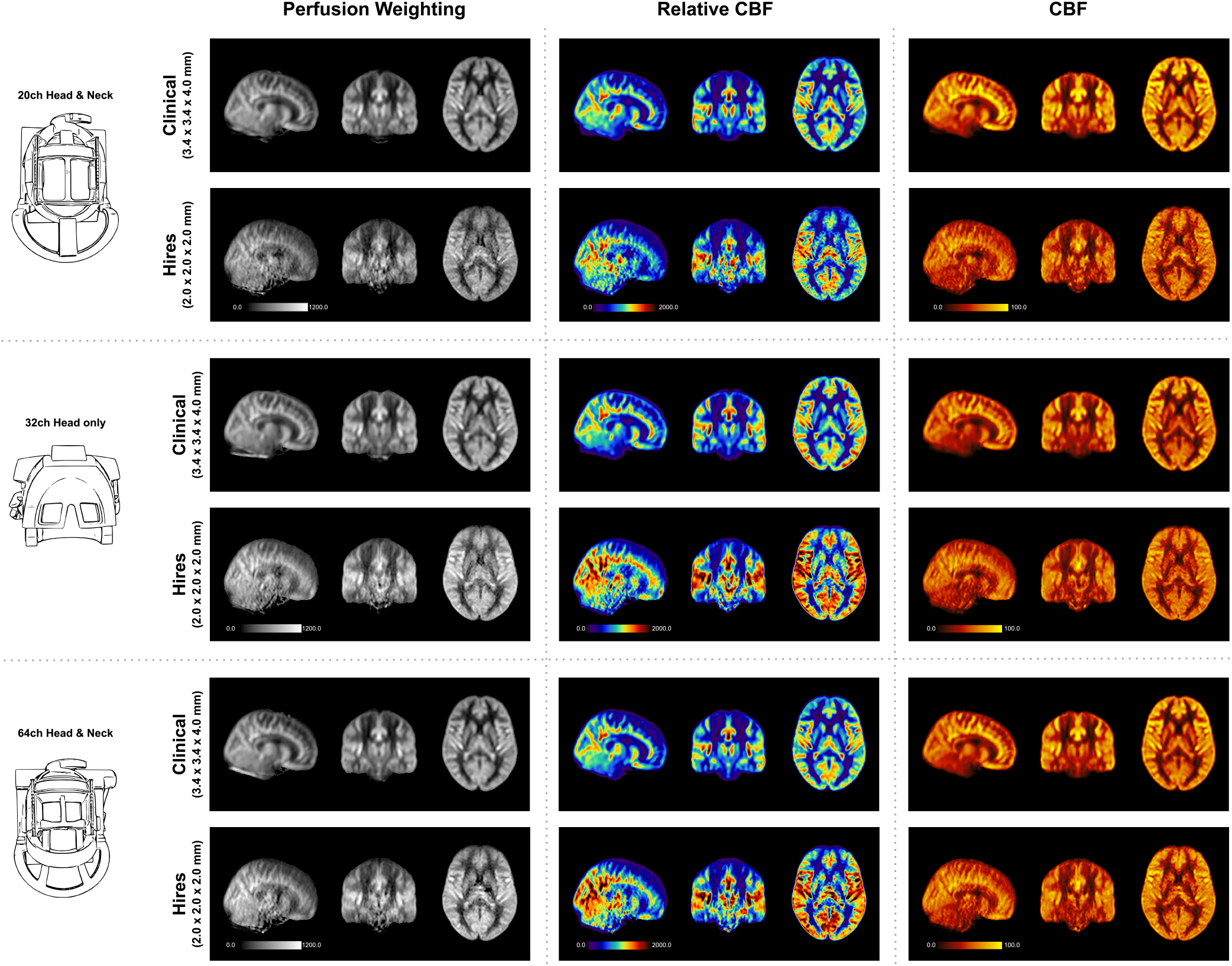
Mosaic of orthogonal views of the group average perfusion-weighted (MRI signal units), relative CBF (arbitrary units) and CBF (mL/100g/min) maps for data acquired using the three respective head coils.. In each panel, the maps obtained from the clinical and hires pCASL acquisitions are displayed in the top and bottom rows, respectively.

#### Tables of Summary Statistics : Clinical vs Hires pCASL

**Table S2.** Summary Statistics of quantitative CBF (in mL/100 g/min) for Clinical and Hires pCASL data over all subjects in the study.

| CBF (mL/100 g/min)<br>pCASL Clinical |  |  |  | CBF (mL/100 g/min)<br>pCASL Hires |  |  |  |
| --- | --- | --- | --- | --- | --- | --- | --- |
| Statistic | 20 ch | 32 ch | 64 ch | Statistic | 20 ch | 32 ch | 64 ch |
| Mean | 72.57 | 68.43 | 67.36 | Mean | 57.90 | 58.30 | 59.56 |
| Median | 70.81 | 65.42 | 64.42 | Median | 56.55 | 56.66 | 57.75 |
| Standard Deviation | 27.85 | 27.96 | 27.61 | Standard Deviation | 26.15 | 25.07 | 25.77 |
| Inter-quartile Range | 36.85 | 37.70 | 37.12 | Inter-quartile Range | 35.58 | 33.59 | 34.85 |
| 1st Quartile | 53.55 | 48.65 | 47.79 | 1st Quartile | 39.42 | 40.83 | 41.44 |
| 3rd Quartile | 90.40 | 86.35 | 84.91 | 3rd Quartile | 75.00 | 74.42 | 76.28 |
| Outlier Percentage | 0.79% | 0.80% | 0.88% | Outlier Percentage | 0.69% | 0.78% | 0.74% |

**Table S3.** Summary Statistics of relative CBF for Clinical and Hires pCASL data over all subjects in the study.

| Relative CBF<br>pCASL Clinical |  |  |  | Relative CBF<br>pCASL Hires |  |  |  |
| --- | --- | --- | --- | --- | --- | --- | --- |
| Statistic | 20 ch | 32 ch | 64 ch | Statistic | 20 ch | 32 ch | 64 ch |
| Mean | 960.79 | 1016.20 | 1001.84 | Mean | 944.67 | 1072.07 | 1097.49 |
| Median | 933.14 | 963.99 | 950.55 | Median | 885.92 | 1003.30 | 1023.64 |
| Standard Deviation | 385.91 | 435.08 | 433.61 | Standard Deviation | 502.92 | 561.64 | 578.83 |
| Inter-quartile Range | 533.11 | 566.97 | 574.38 | Inter-quartile Range | 693.79 | 759.00 | 788.37 |
| 1st Quartile | 678.91 | 704.31 | 687.94 | 1st Quartile | 568.31 | 656.43 | 666.43 |
| 3rd Quartile | 1212.03 | 1271.28 | 1262.31 | 3rd Quartile | 1262.09 | 1415.43 | 1454.80 |
| Outlier Percentage | 0.81% | 1.66% | 1.46% | Outlier Percentage | 0.99% | 1.28% | 1.18% |

**Table S4.** Summary Statistics of Perfusion-weighting (in arbitrary units) for Clinical and Hires pCASL data over all subjects in the study.

| Perfusion weighting (a. u.)<br>pCASL Clinical |  |  |  | Perfusion weighting (a. u.)<br>pCASL Hires |  |  |  |
| --- | --- | --- | --- | --- | --- | --- | --- |
| Statistic | 20 ch | 32 ch | 64 ch | Statistic | 20 ch | 32 ch | 64 ch |
| Mean | 575.50 | 608.55 | 600.29 | Mean | 534.89 | 605.91 | 620.15 |
| Median | 557.94 | 576.44 | 568.68 | Median | 500.93 | 565.98 | 577.39 |
| Standard Deviation | 226.10 | 256.24 | 256.66 | Standard Deviation | 280.25 | 313.30 | 322.90 |
| Inter-quartile Range | 312.50 | 334.30 | 338.79 | Inter-quartile Range | 386.34 | 423.67 | 440.03 |
| 1st Quartile | 409.56 | 424.01 | 414.27 | 1st Quartile | 324.63 | 373.16 | 378.73 |
| 3rd Quartile | 722.06 | 758.30 | 753.05 | 3rd Quartile | 710.97 | 796.82 | 818.77 |
| Outlier Percentage | 0.84% | 1.69% | 1.51% | Outlier Percentage | 1.03% | 1.31% | 1.21% |

**Table S5.** Summary Statistics of perfusion tSNR for Clinical and Hires pCASL data over all subjects in the study.

| Perfusion tSNR<br>pCASL Clinical |  |  |  | Perfusion tSNR<br>pCASL Hires |  |  |  |
| --- | --- | --- | --- | --- | --- | --- | --- |
| Statistic | 20 ch | 32 ch | 64 ch | Statistic | 20 ch | 32 ch | 64 ch |
| Mean | 5.66 | 5.77 | 5.92 | Mean | 1.59 | 2.14 | 2.27 |
| Median | 5.43 | 5.51 | 5.64 | Median | 1.45 | 1.85 | 1.97 |
| Standard Deviation | 2.35 | 2.45 | 2.53 | Standard Deviation | 0.92 | 1.39 | 1.44 |
| Inter-quartile Range | 3.06 | 3.17 | 3.23 | Inter-quartile Range | 1.16 | 1.63 | 1.74 |
| 1st Quartile | 3.99 | 4.03 | 4.14 | 1st Quartile | 0.93 | 1.16 | 1.24 |
| 3rd Quartile | 7.06 | 7.20 | 7.37 | 3rd Quartile | 2.09 | 2.79 | 2.98 |
| Outlier Percentage | 1.50% | 1.67% | 1.83% | Outlier Percentage | 2.29% | 3.55% | 3.20% |

**Table S6.** Summary Statistics of perfusion tSNR after scaling the pCASL hires data in Table S7 by the ratio of voxel-volumes.

| Perfusion tSNR<br>pCASL Clinical |  |  |  | Perfusion tSNR (volume-scaled)<br>pCASL Hires |  |  |  |
| --- | --- | --- | --- | --- | --- | --- | --- |
| Statistic | 20 ch | 32 ch | 64 ch | Statistic | 20 ch | 32 ch | 64 ch |
| Mean | 5.66 | 5.77 | 5.92 | Mean | 9.19 | 12.35 | 13.11 |
| Median | 5.43 | 5.51 | 5.64 | Median | 8.40 | 10.66 | 11.40 |
| Standard Deviation | 2.35 | 2.45 | 2.53 | Standard Deviation | 5.30 | 8.03 | 8.34 |
| Inter-quartile Range | 3.06 | 3.17 | 3.23 | Inter-quartile Range | 6.70 | 9.42 | 10.03 |
| 1st Quartile | 3.99 | 4.03 | 4.14 | 1st Quartile | 5.38 | 6.70 | 7.18 |
| 3rd Quartile | 7.06 | 7.20 | 7.37 | 3rd Quartile | 12.08 | 16.12 | 17.21 |
| Outlier Percentage | 1.50% | 1.67% | 1.83% | Outlier Percentage | 2.29% | 3.55% | 3.20% |

### Comparison of Hires pCASL and PASL protocols

**Figure S2.**
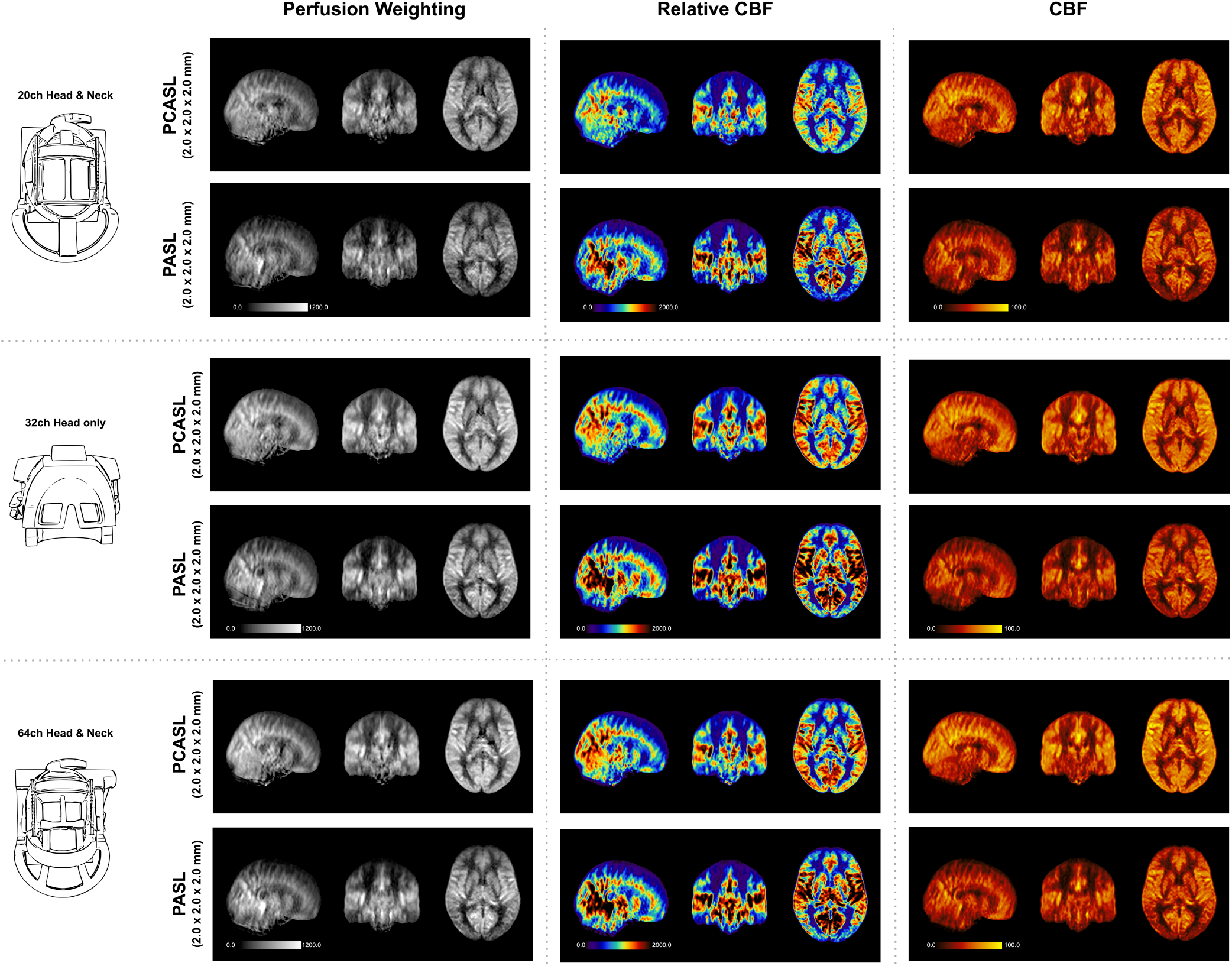
Mosaic of orthogonal views of the group average perfusion-weighted (MRI signal units), relative CBF (arbitrary units) and CBF (mL/100g/min) maps for data acquired using the three respective head coils. In each panel, the maps obtained from the hires pCASL and PASL acquisitions are displayed in the top and bottom rows, respectively.

#### Impact of head coil choice on hires acquisitions

**Figure S3.**
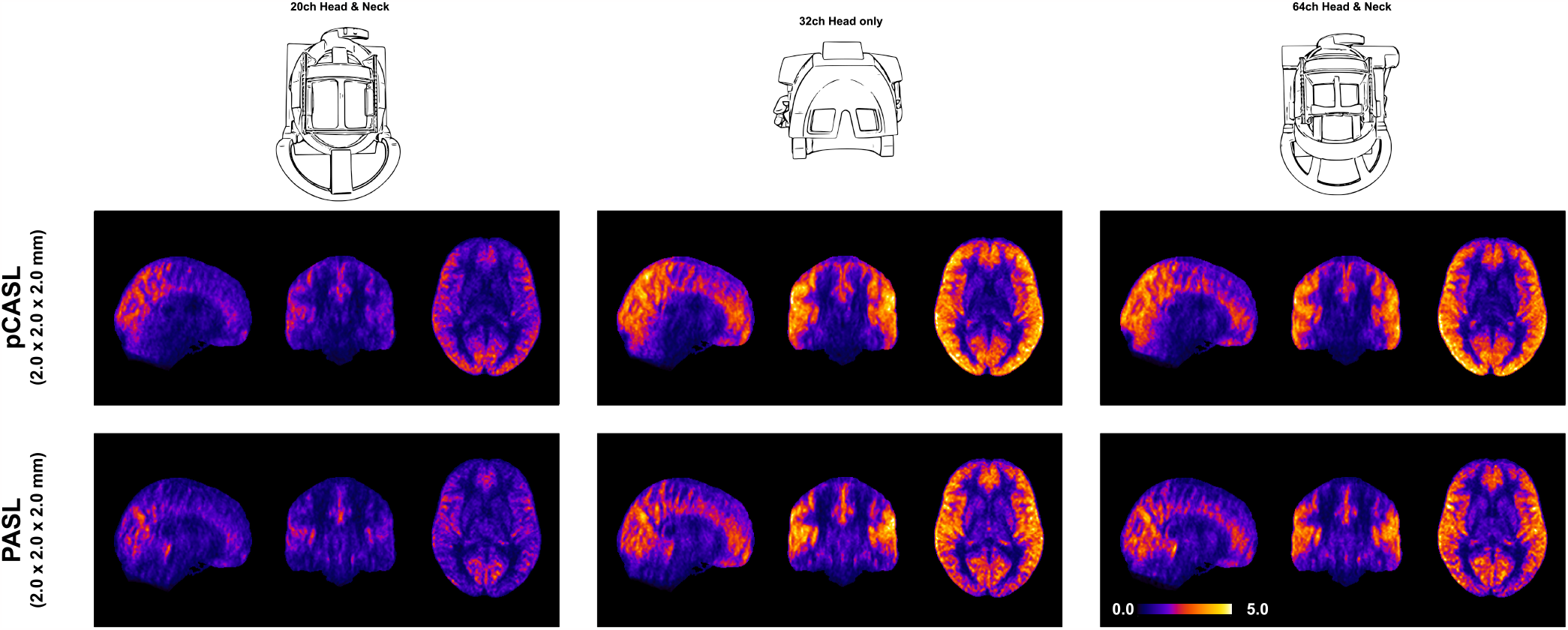
Orthogonal views of the group average perfusion tSNR maps for the hires acquisitions with pCASL and PASL labelling schemes.

#### Tables of Summary Statistics : Hires pCASL vs Hires PASL

**Table S7.** Summary Statistics of quantitative CBF (in mL/100 g/min) for pCASL Hires and PASL Hires data over all subjects in the study.

| CBF (mL/100 g/min)<br>pCASL Hires |  |  |  | CBF (mL/100 g/min)<br>PASL Hires |  |  |  |
| --- | --- | --- | --- | --- | --- | --- | --- |
| Statistic | 20 ch | 32 ch | 64 ch | Statistic | 20 ch | 32 ch | 64 ch |
| Mean | 57.90 | 58.30 | 59.56 | Mean | 42.06 | 42.12 | 42.37 |
| Median | 56.55 | 56.66 | 57.75 | Median | 39.65 | 39.82 | 40.20 |
| Standard Deviation | 26.15 | 25.07 | 25.77 | Standard Deviation | 25.94 | 25.71 | 25.80 |
| Inter-quartile Range | 35.58 | 33.59 | 34.85 | Inter-quartile Range | 37.10 | 36.93 | 37.09 |
| 1st Quartile | 39.42 | 40.83 | 41.44 | 1st Quartile | 21.86 | 22.11 | 22.30 |
| 3rd Quartile | 75.00 | 74.42 | 76.28 | 3rd Quartile | 58.96 | 59.04 | 59.39 |
| Outlier Percentage | 0.69% | 0.78% | 0.74% | Outlier Percentage | 0.77% | 0.72% | 0.71% |

**Table S8.** Summary Statistics of relative CBF for pCASL Hires and PASL Hires data over all subjects in the study.

| Relative CBF<br>pCASL Hires |  |  |  | Relative CBF<br>PASL Hires |  |  |  |
| --- | --- | --- | --- | --- | --- | --- | --- |
| Statistic | 20 ch | 32 ch | 64 ch | Statistic | 20 ch | 32 ch | 64 ch |
| Mean | 944.67 | 1072.07 | 1097.49 | Mean | 988.63 | 1114.79 | 1126.50 |
| Median | 885.92 | 1003.30 | 1023.64 | Median | 863.30 | 970.60 | 984.09 |
| Standard Deviation | 502.92 | 561.64 | 578.83 | Standard Deviation | 701.69 | 793.27 | 801.86 |
| Inter-quartile Range | 693.79 | 759.00 | 788.37 | Inter-quartile Range | 978.52 | 1103.48 | 1121.65 |
| 1st Quartile | 568.31 | 656.43 | 666.43 | 1st Quartile | 435.20 | 489.73 | 492.45 |
| 3rd Quartile | 1262.09 | 1415.43 | 1454.80 | 3rd Quartile | 1413.71 | 1593.21 | 1614.10 |
| Outlier Percentage | 0.99% | 1.28% | 1.18% | Outlier Percentage | 1.32% | 1.38% | 1.30% |

**Table S9.** Summary Statistics of Perfusion-weighting (in arbitrary units) for pCASL Hires and PASL Hires data over all subjects in the study.

| Perfusion weighting (a. u.)<br>pCASL Hires |  |  |  | Perfusion weighting (a. u.)<br>PASL Hires |  |  |  |
| --- | --- | --- | --- | --- | --- | --- | --- |
| Statistic | 20 ch | 32 ch | 64 ch | Statistic | 20 ch | 32 ch | 64 ch |
| Mean | 534.89 | 605.91 | 620.15 | Mean | 420.22 | 471.74 | 476.20 |
| Median | 500.93 | 565.98 | 577.39 | Median | 369.42 | 412.66 | 417.72 |
| Standard Deviation | 280.25 | 313.30 | 322.90 | Standard Deviation | 286.37 | 324.00 | 327.51 |
| Inter-quartile Range | 386.34 | 423.67 | 440.03 | Inter-quartile Range | 396.12 | 447.66 | 454.97 |
| 1st Quartile | 324.63 | 373.16 | 378.73 | 1st Quartile | 196.43 | 218.29 | 219.09 |
| 3rd Quartile | 710.97 | 796.82 | 818.77 | 3rd Quartile | 592.56 | 665.95 | 674.06 |
| Outlier Percentage | 1.03% | 1.31% | 1.21% | Outlier Percentage | 1.38% | 1.44% | 1.36% |

**Table S10.**
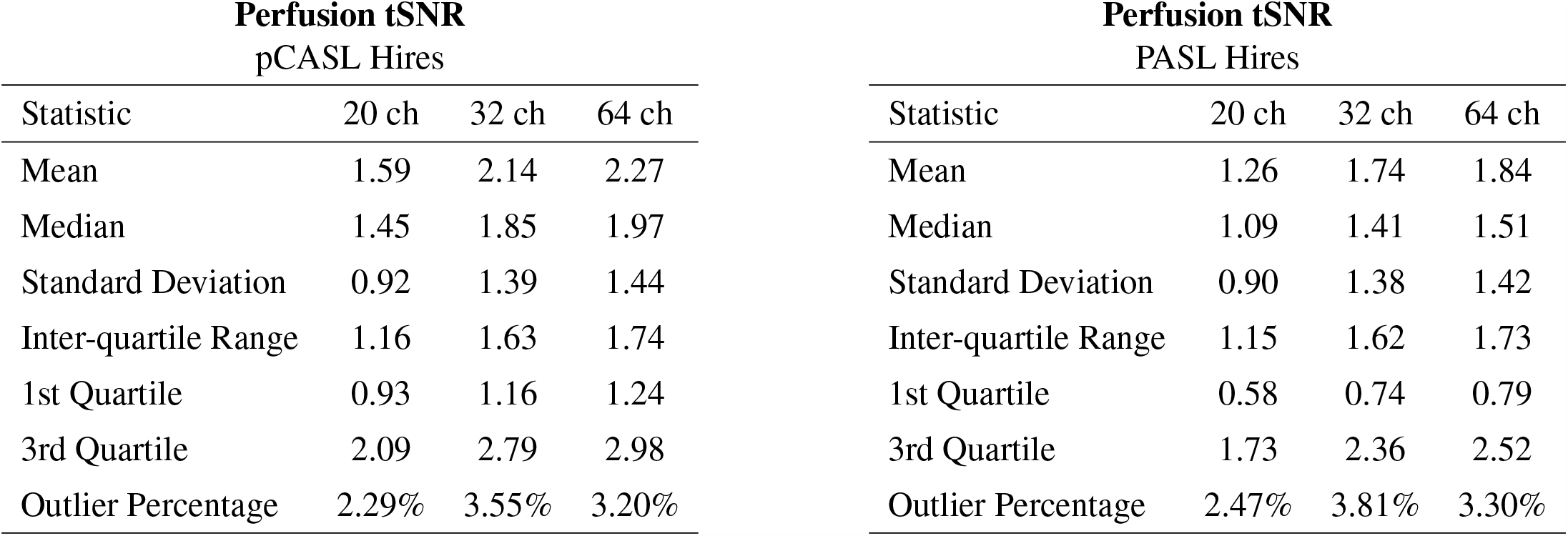
Summary Statistics of perfusion tSNR for pCASL Hires and PASL Hires data over all subjects in the study.

**Table S11.** Summary Statistics of perfusion tSNR after scaling the pCASL hires data in Table S7 by the ratio of voxel-volumes.

| Perfusion tSNR (volume-scaled)<br>pCASL Hires |  |  |  |
| --- | --- | --- | --- |
| Statistic | 20 ch | 32 ch | 64 ch |
| Mean | 9.19 | 12.35 | 13.11 |
| Median | 8.40 | 10.66 | 11.40 |
| Standard Deviation | 5.30 | 8.03 | 8.34 |
| Inter-quartile Range | 6.70 | 9.42 | 10.03 |
| 1st Quartile | 5.38 | 6.70 | 7.18 |
| 3rd Quartile | 12.08 | 16.12 | 17.21 |
| Outlier Percentage | 2.29% | 3.55% | 3.20% |

| Perfusion tSNR (volume-scaled)<br>PASL Hires |  |  |  |
| --- | --- | --- | --- |
| Statistic | 20 ch | 32 ch | 64 ch |
| Mean | 7.28 | 10.06 | 10.63 |
| Median | 6.28 | 8.12 | 8.72 |
| Standard Deviation | 5.18 | 8.00 | 8.24 |
| Inter-quartile Range | 6.63 | 9.39 | 10.02 |
| 1st Quartile | 3.37 | 4.25 | 4.54 |
| 3rd Quartile | 10.00 | 13.64 | 14.56 |
| Outlier Percentage | 2.47% | 3.81% | 3.30% |

### Single participant maps for all three ASL protocols

**Figure S4.**
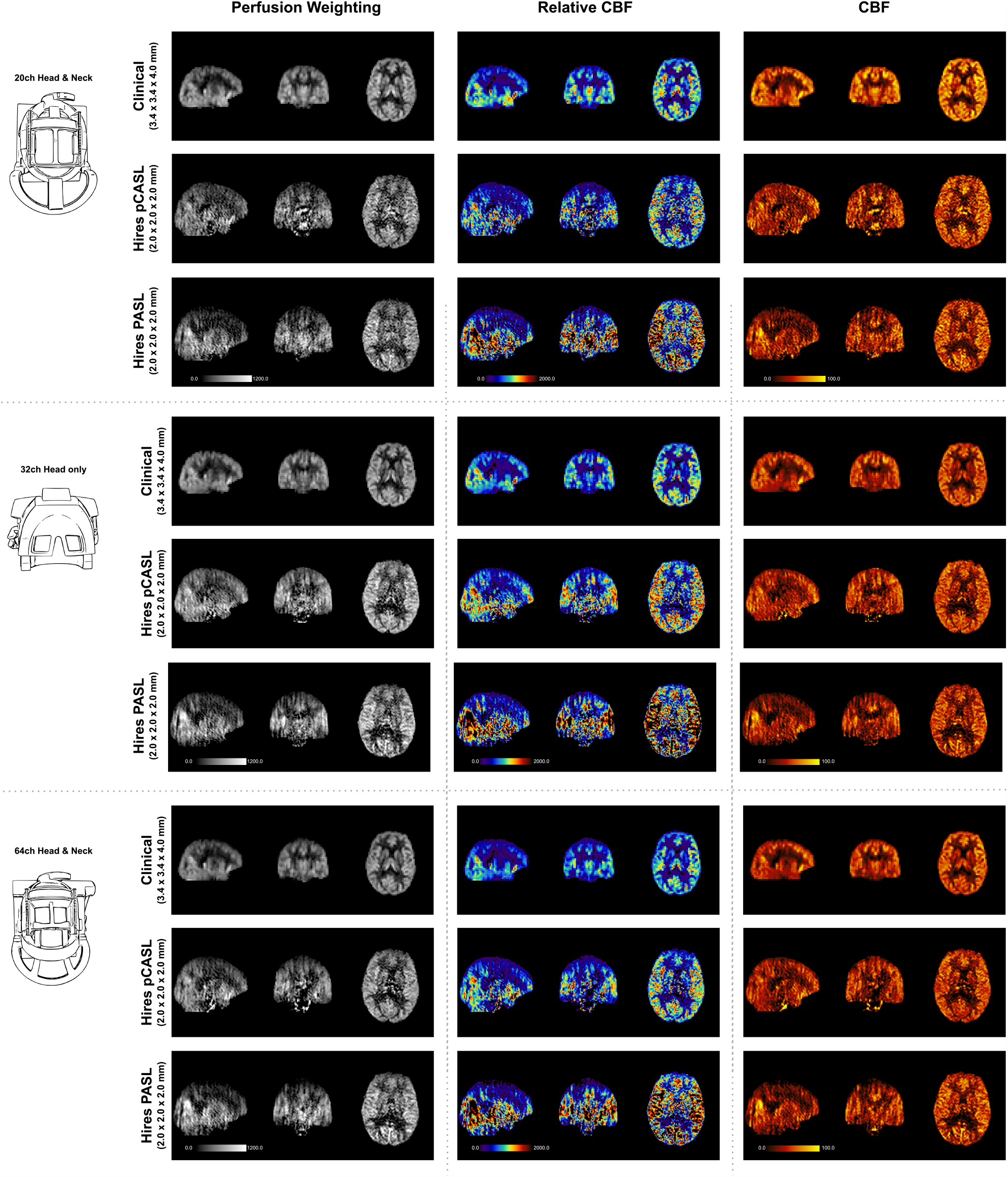
Mosaic of orthogonal views of a single participant’s perfusion-weighted (MRI signal units), relative CBF (arbitrary units) and CBF (mL/100g/min) maps for data acquired using the three respective head coils. In each panel, the maps obtained from the clinical pCASL, hires pCASL and hires PASL acquisitions are displayed in the top, middle and bottom rows, respectively.

